# Genomic evidence for atmospheric chemosynthesis as a defining trait of Ktedonobacteria inhabiting subterranean environments

**DOI:** 10.64898/2025.12.07.692797

**Authors:** Andrea Firrincieli, Giacomo Broglia, Giulia Rizzo, Daniele Ghezzi, Francesco Sauro, Martina Cappelletti

## Abstract

Members of the class Ktedonobacteria (phylum Chloroflexota) are widespread across a variety of terrestrial environments. They are particularly enriched in silica-rich oligotrophic caves, where they may act as early colonizers, but the genomic bases supporting this pioneering capacity are still unclear. Here, we present a systematic genomic analysis of Ktedonobacteria from diverse ecosystems, with a focus on oligotrophic caves. Cave Ktedonobacteria belong to novel genera of *Ktedonobacteraceae* and harbor genes supporting a mixotrophic metabolism that combines the use of organic and inorganic substrates as energy and carbon sources. Comparative analyses of all available Ktedonobacteria genomes from diverse environments showed that most metabolic traits, including those enabling atmospheric gas oxidation, are primarily driven by taxonomy. In contrast, genes involved in CO_2_ fixation are enriched in Ktedonobacteria inhabiting caves. Phylogenetic analysis showed that the RuBisCO of Ktedonobacteria likely represents a novel Form I subtype, which we designate as BAK, reflecting its wide occurrence in thermophilic and acidophilic Bacillota and Actinomycetota inhabiting similar extreme environments. The presence of this subtype across distinct lineages in comparable habitats suggests that it may confer a selective advantage in nutrient-poor settings (like caves) and that its distribution may be subject to horizontal gene transfer. This autotrophic capacity relies on a transaldolase variant of the Calvin–Benson–Bassham cycle that was previously described only in one Firmicutes species. Overall, this study provides genetic evidence supporting the combination of atmospheric gas oxidation with dark CO_2_ fixation in Ktedonobacteria, highlighting their role as primary producers in oligotrophic ecosystems, including caves.

## Introduction

Caves are subterranean environments that support rich and diverse microbial communities, despite being generally characterized by oligotrophic conditions (i.e., low nutrient availability) in association to the sporadic access of organic compounds from the surface and the absence of sunlight-driven photosynthesis [1–3]. This is because cave microorganisms have evolved metabolic strategies based on chemolithotrophic processes and the ability to establish synergistic interactions involving metabolite exchange [4, 5]. Among these strategies, cave microbes can scavenge nutrients released through rock weathering and from the oxidation of atmospheric trace gases (H_2_, CO, and CH_4_) [6, 7]. A metagenomic study further revealed an overrepresentation of RuBisCO genes in a speleothem sample from a carbonate cave compared with metagenomes from soil, rhizosphere, and marine environments, suggesting that CO_2_ fixation in caves may occur through metabolic strategies independent of sunlight energy [8]. Quartzite caves of the Guiana Shield tepuis (among the oldest geological formations on Earth) offer exceptional natural laboratories to investigate microbial colonization and rock–microbe interactions under aphotic and oligotrophic conditions. These ancient and remote subterranean systems are characterized by stable temperature and humidity, slight acidity due to the limited buffering capacity of quartzite, and an almost complete absence of organic nutrients [9, 10]. Previous studies have revealed complex and diverse microbial communities whose composition is strongly influenced by silica dissolution/precipitation processes and water availability [9, 11, 12]. Microbiological analyses further indicated that early colonization stages are dominated by members of the class Ktedonobacteria that are unclassified at lower taxonomy levels [6, 11].

Ktedonobacteria (phylum *Chloroflexota*) are mostly filamentous, aerobic bacteria with chemoheterotrophic capacities when considering the cultivated isolates described so far. They have been predominantly isolated and characterized from soil environments, although DNA-based surveys consistently report their presence in nutrient-poor and extreme habitats, including volcanic soils, geothermal fields, deserts, and silica-rich caves, like quartzite caves and the fumarolic ice cave Warren (Mt. Erebus, Antarctica) [13–15]. Targeted gene analyses revealed that most of the genetic diversity associated with the atmospheric gas oxidation and CO_2_ fixation in both cave systems derived from members of Ktedonobacteria [6, 16]. Furthermore, several studies indicated these bacteria to dominate the early stages of microbial colonisation processes in both volcanic soil and in oligotrophic quartzite caves [6, 9, 17]. Despite their occurrence in oligotrophic caves and their possible key role as pioneer colonisers of subterranean environments, a comparative genomic investigation of the metabolic features that distinguish cave-associated Ktedonobacteria from those inhabiting other ecosystems is still lacking.

In this work, we present a systematic genomic investigation of Ktedonobacteria thriving in diverse ecological niches, including aphotic oligotrophic subterranean environments (caves). By classifying genomes according to their ecosystem of origin and performing comparative analyses, we identified genetic traits linked to cave colonisation, providing insights into the metabolic strategies that may support the early colonisation and persistence of *Ktedonobacteria* in dark, oligotrophic ecosystems.

## Materials and Methods

### DNA extraction and shotgun metagenomic sequencing

Total microbial DNA extractions (of the samples Ay304, Ay317 and Ay302 from Imawarì Yeutà) were performed using the PowerSoil Lyzer Kit (Qiagen) as previously described [12]. Fifty ng of DNA were used as input material for the sequencing library preparation using the NEBNext Ultra™ II FS DNA Kit (New England Biolabs). The library was sequenced on a NextSeq 500 (Illumina) with the following specifications: read length 150 bp, paired-end, 7.5 Gb per sample. The DNA obtained from Ay302 and Ay317 samples was also sequenced using Oxford Nanopore Technology (ONT). ONT sequencing libraries were prepared and sequenced using the SQK-LSK110 kit (ONT). The sequencing libraries were loaded on a FLO-MIN106D flow cell (chemistry R9.4.1) and sequenced on a MinION Mk1C device. Due to insufficient amount of DNA and starting material, the sample Ay304 was only sequenced through Illumina, but not by ONT.

### Metagenome assembly and reconstruction of metagenome-assembled genomes (MAGs)

For the reconstruction of MAGs from Illumina sequencing data obtained from the fumarolic ice cave Warren (Mt. Erebus, PRJNA255918) and the Ay304 sample, raw reads were subjected to adapter removal and quality trimming using bbduk (https://sourceforge.net/projects/bbmap/), and finally assembled with MEGAHIT [18]. Contigs longer than 2.5 Kbp were binned using the metaWRAP ‘binning’ and ‘binning_refinement’ modules with default parameters [19]. The resulting bins with completeness > 50% and contamination < 10% were taxonomically classified using the Genome Taxonomy Database (GTDB, http://gtdb.ecogenomic.org) (GTDB-Tk) v.2.4.1 [20]. The quality of MAGs classified as Ktedonobacteria was further assessed using BUSCO v5.4.4 [21].

In the case of Ay317 and Ay302 metagenomes (sequenced via both Illumina and ONT), MAG reconstruction was performed using the MetaPlatanus [22] pipeline in combination with OPERA-MS [23] and metaWRAP. First, the ONT ‘*fast5’* signal data were basecalled with gpu-guppy v6.2.0 using the super-accurate basecalling configuration (dna_r9.4.1_450bps_sup.cfg) with the following parameters enabled: ‘*--detect_adapter*’, ‘*--detect_mid_strand_adapter*’, and ‘*--trim_adapters*’. High-quality Illumina reads were obtained as described above, using bbduk for adapter removal and quality trimming. High-quality ONT and Illumina reads were subsequently used as input for MetaPlatanus. The resulting scaffolds obtained after the gap-closing and polishing steps with TGS-GapCloser and NextPolish, respectively, were binned using the OPERA-MS binning module ‘*OPERA-MS-UTILS*.*py binning*’ and finally refined with ‘binning_refinement’ module of metaWRAP. The resulting bins were evaluated for completeness and contamination, taxonomically classified using GTDB-Tk v2.4.1, and quality-assessed with BUSCO v5.4.4 for those identified as Ktedonobacteria.

### Functional annotation, phylogenomic and comparative analyses

Functional annotation and metabolic potential prediction of Ktedonobacteria MAGs recovered from Imawarì Yeutà, Warren [16], and Monte Cristo caves [24] were performed using the Anvi’o v7.1 contigs workflow using both ‘anvi_run_kegg_kofams’ and ‘anvi_run_pfams’ against the KEGG and InterPro functional databases, respectively [25]. In addition, dbCAN v3 was used to predict the carbohydrate degradation potential [26]. A phylogenomic tree of the Ktedonobacteria was generated using the GTDB-Tk de novo workflow. The concatenated alignment of 120 single-copy bacterial marker genes (bac120) was used as input for IQTREE v2.2.5 [27] to compute a maximum-likelihood phylogeny using the WAG substitution model and ultrafast bootstrap (1,000 replicates). The resulting tree was visualised and annotated using iTOL [28].

The pangenomic analysis of Ktedonobacteria MAGs and genomes was performed using the Anvi’o v7.1 [25] pangenomic workflow. The input files for the Anvi’o pangenome analysis were represented by the genome contig files recovered from NCBI GenBank database, and from this study (Table S1). Within the Anvi’o pangenome workflow, the metabolic potential of each Ktedonobacteria was carried out by first identifying open reading frames with the anvi_gen_contigs_database script, followed by functional assignment of the resulting amino acid sequences using anvi_run_kegg_kofams and anvi_run_pfams against the KEGG and InterPro databases, respectively. Finally, pangenomic analysis was performed using the anvi_pan_genome script with the ‘mcl-inflation’ parameter set to 2, to identify clusters of ortholog genes between distantly related genomes.

### Phylogenetic and synteny analyses of RuBisCO genes and gene clusters in Ktedonobacteria

Non-redundant (90% cluster identity) prokaryotic RuBisCO large subunit (*rbcL*) sequences, representing different subtypes of the Forms I, II, III, and IV, were retrieved from Prywes *et al*. [24]. Sequences were aligned with MAFFT v7.310 [25] using the G-INS-i algorithm, and poorly aligned regions were trimmed with trimAl in automated mode [26]. The alignment file was analysed in IQTREE v2.2.5 [27] to build a maximum-likelihood phylogenetic tree with the following settings: ultra-fastbootstrap (-bb) ‘1000’, model-finder (-m) ‘TEST’, and threads (-T) ‘AUTO’. The resulting consensus tree was constructed under the LG+G4 substitution model and uploaded in iTOL for visualisation (https://itol.embl.de/shared/m11l1Qm6EqfZ). Synteny analysis of RuBisCO gene clusters in Ktedonobacteria was performed using clinker with default parameters (https://github.com/gamcil/clinker). Briefly, contigs possessing one copy each RuBisCO subunit were reannotated using prokka 1.14.6 [28]. The resulting GenBank files were finally aligned with clinker to visualise gene cluster organisation and conservation (https://github.com/gamcil/clinker).

### Statistical analyses

Jaccard distances were computed from *Ktedonobacteria* KEGG Ortholog (KO) presence/absence matrices generated with Anvi’o, using the microViz R package v0.12.4. A PERMANOVA with 999 permutations was then conducted to assess differences in KO composition among Ktedonobacteria families. The identification of family- and SE-specific features was performed using the Rao’s Score test implemented in the script ‘anvi-compute-functional-enrichment-in-pan’. The analysis was performed to identify both family-specific and niche-specific functional genes according to an adjusted q-value < 0.001 and < 0.01, respectively.

## Results

### Reconstruction of Ktedonobacteria genomes from oligotrophic caves

Metagenome-assembled genomes (MAGs) of Ktedonobacteria inhabiting cave environments were reconstructed from metagenomic sequencing data obtained from samples collected in the Imawarì Yeutà cave (Auyán Tepui, Venezuela), Monte Cristo cave (Diamantina, Brazil), and Warren Cave (Mt. Erebus, Antarctica). These metagenomes were selected because previous 16S rRNA gene analyses revealed detectable presence of members of this class (Table S2).

Metagenomic assemblies from the Imawarì Yeutà cave generated four MAGs affiliated with the class Ktedonobacteria. Specifically, the metagenomic assembly from sample Ay304 yielded only one MAG assigned to Ktedonobacteria. In contrast, the assembly from sample Ay302 produced several MAGs, three of which belonged to this class (Table S1). Sample Ay317 did not yield any Ktedonobacteria MAGs of sufficient quality. The metagenomic assembly of datasets from Warren Cave yielded two MAGs affiliated with Ktedonobacteria (Table S1). Together with the previously reconstructed MAGs from the Monte Cristo cave [29], these genomes compose the dataset of eight Ktedonobacteria MAGs inhabiting oligotrophic caves, which were subsequently analyzed for their metabolic potential.

### The metabolic potential of cave Ktedonobacteria

#### Central carbon metabolism and carbon fixation

Ktedonobacteria MAGs from oligotrophic caves possess genes coding for enzymes of complete/nearly complete pathways of glycolysis/gluconeogenesis, pentose phosphate cycle, tricarboxylic (TCA) cycle, synthesis and β-oxidation of fatty acids, CO_2_ fixation through the Calvin–Benson–Bassham (CBB) cycle (Fig. 1).

**Figure 1.**
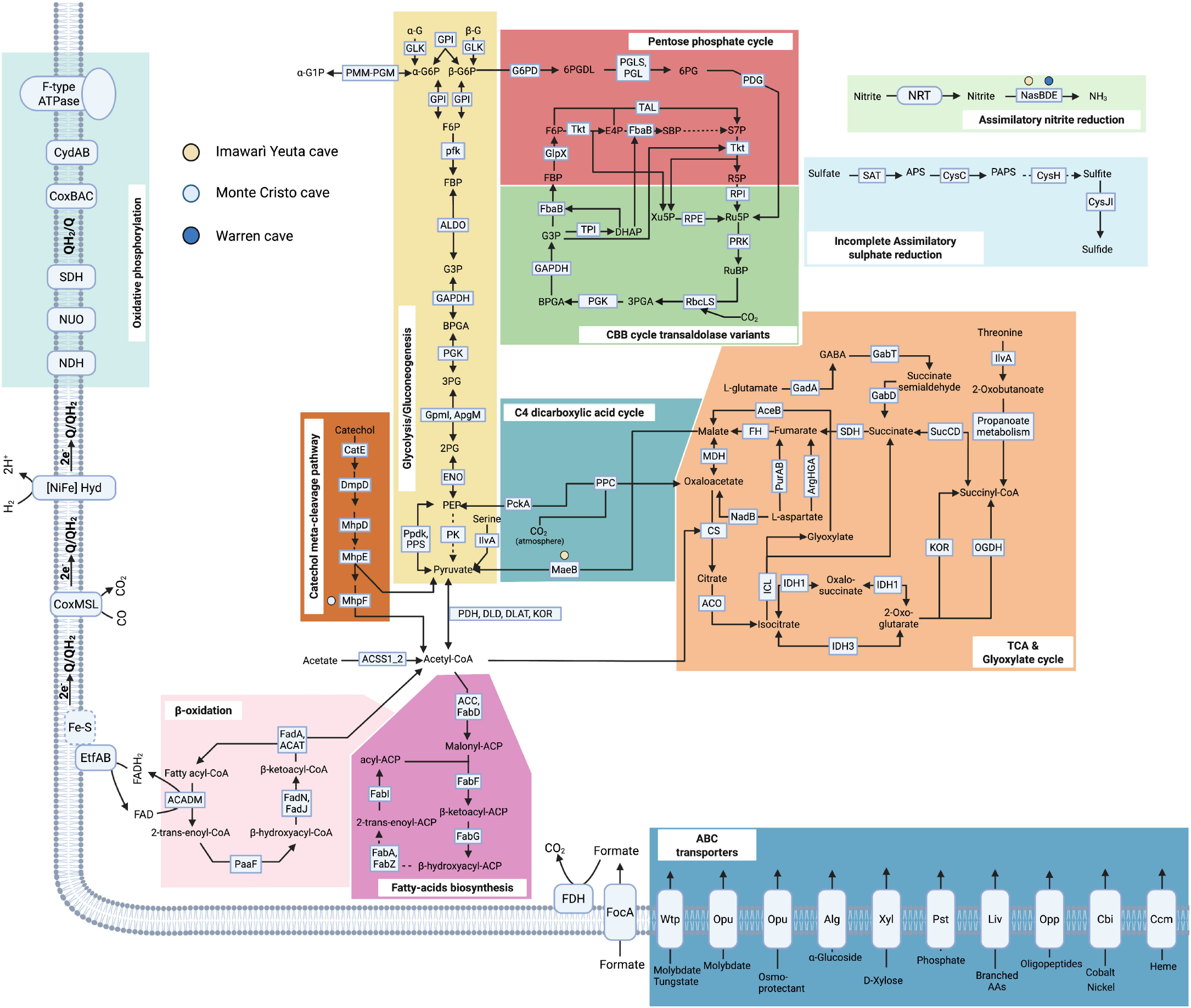
Genome-based metabolic reconstruction of Ktedonobacteria inhabiting silica-rich oligotrophic caves. Coloured dots indicate reactions uniquely associated with MAGs from caves. A summary of the functional annotation for all MAGs used to generate this figure is provided in Table S3. Abbreviations of metabolites: α/β-G, α/β-glucose; α/β-G6P, α/β-glucose-6-phosphate; α-G1P, α-glucose-1-phosphate; 3PG, 3-phosphoglycerate; 2-PG, 2-phosphoglycerate; PEP, phosphoenolpyruvate; GABA, γ-aminobutyric acid; RuBP, ribulose-1,5-bisphosphate; 3PGA, 3-phosphoglycerate; BPGA, 1,3-bisphosphoglycerate; DHAP, dihydroxyacetone phosphate; G3P, glyceraldehyde-3-phosphate; FBP, fructose-1,6-bisphosphate; F6P, fructose-6-phosphate; E4P, erythrose-4-phosphate; SBP, sedoheptulose-1,7-bisphosphate; S7P, sedoheptulose-7-phosphate; R5P, ribose-5-phosphate; Ru5P, ribulose-5-phosphate.

Analysis of the glycolytic pathway indicates that all cave Ktedonobacteria possess the genes involved in Embden–Meyerhof–Parnas (EMP) variant of glycolysis. However, MAGs from Imawarì Yeutà lack the pyruvate kinase (*pk*, K00873) gene, an absence apparently compensated by the presence of the PPi-dependent pyruvate, orthophosphate dikinase (*ppdk*, K01006) and pyruvate, water dikinase (*pps*, K01007) genes, both of which catalyse the reversible conversion of phosphoenolpyruvate (PEP) to pyruvate. Cave Ktedonobacteria also possess a complete set of genes involved in the oxidative (module M00006) and non-oxidative (module M00007) phases of the pentose phosphate pathway allowing to generate precursors for the biosynthesis of nucleotides and amino acids. Ktedonobacteria from Monte Cristo cave possess the genes of the catechol meta-cleavage pathway.

A complete set of genes encoding the enzymes of the TCA cycle was detected, together with additional genes catalyzing anaplerotic reactions that replenish TCA intermediates. Specifically, cave *Ktedonobacteria* possess i) the gene encoding phosphoenolpyruvate carboxylase (*ppc*, K01595), which catalyses the addition of CO2 to phosphoenolpyruvate to form oxaloacetate, ii) the genes encoding isocitrate lyase (*icl*, K01637) and malate synthase (*aceB*, K01638), two key enzymes of the glyoxylate cycle that convert two acetyl-CoA molecules into succinate. The resulting succinate can subsequently enter the TCA cycle or be used for amino acid biosynthesis. Other genes indicate the capability of cave *Ktedonobacteria* to use amino acids as carbon sources through the synthesis of various TCA intermediates. These include the formation of fumarate via the activity of enzymes encoded by *purAB* (K01939-K01756) and *argGHA* (K01940-K01755-K1468); oxalacetate via aspartate deamination (*nadB*, K00278), succinate via glutamate conversion (*gadA*, K01580; *gabT*, K07250; *gabD*, K00135); and succinyl-CoA via threonine deamination in the propanoyl-CoA metabolism (*ilvA*, K01754 in module M00741).

With respect to carbon fixation, cave Ktedonobacteria lack genes encoding the key enzymes of the 3-hydroxypropionate bicycle and the Wood-Ljungdahl pathway, which are typically present in anaerobic and facultative aerobic members of Chloroflexota performing light-driven photosynthesis [15]. Conversely, cave Ktedonobacteria possess genes encoding the large (*rbcL*, K01601) and small (*rbcS*, K01602) subunits of ribulose bisphosphate carboxylase/oxygenase (RuBisCO), as well as the gene encoding transaldolase/fructose-6-phosphate aldolase (TAL, PF00923). The latter enzyme is involved in the transaldolase variant of the Calvin–Benson–Bassham (CBB) cycle, in which the transaldolase reaction regenerates ribulose-1,5-bisphosphate (RuBP), the CO2 acceptor in the cycle [30].

#### Energy metabolism

The electron transport chain of the cave Ktedonobacteria includes genes encoding the main components of the respiratory chain including those for the non-electrogenic NADH:quinone reductase (*ndh*, K03885), NADH:ubiquinone oxidoreductase (*nuo*, module M00144) complex, succinate dehydrogenase (*sdh*, module M00149), cytochrome bd complex (*cydAB*, module M00153), cytochrome aa3bc complex (*coxBAC*, module M00155) and F-type ATP synthase (module M00157) (Fig. 1). Additional genes were also detected that code for complexes able to transfer electrons either directly to the quinone pool of the respiratory chain or indirectly via reducing equivalents (NADH and FADH_2_) generated through the Nuo complex. In particular, the presence of genes coding for the electron transfer flavoproteins (*eft*, K03522-K03521-K00313) and formate dehydrogenase (*fdh*, K00124) suggests that these bacteria can couple the oxidation of fatty acids (beta-oxidation) and CO_2_ with energy metabolism through the production of FADH_2_ and NADH, respectively. Likewise, the presence of genes encoding the aerobic carbon monoxide dehydrogenase complex (*coxMSL, K03519-K03518-K03520*), and the [NiFe]-hydrogenase (*hyd* genes) complex indicates that the cave Ktedonobacteria have the. genetic potential to generate a proton gradient for ATP synthesis through the oxidation of atmospheric trace gases.

#### Nitrogen and sulfur metabolism

Enzymes involved in the dissimilatory sulphur and nitrogen metabolisms are absent in all cave Ktedonobacteria, suggesting that they are unable to couple the reduction of inorganic nitrogen and sulphur compounds with energy metabolism (Fig. 1). Similarly, cave Ktedonobacteria appear to have a limited capacity to assimilate nitrogen and sulphur in the form of nitrate and sulphate, as they lack the genes encoding the enzymes that catalyse the first reduction step of these compounds into nitrite and sulphite, respectively. Conversely, cave Ktedonobacteria possess the genes encoding the sulphite reductase (NADPH) flavoprotein (*cys*) and the nitrite reductase [NAD(P)H] (*nas*), which catalyse the final reactions of the nitrate and sulphate assimilatory pathways, converting nitrite and sulphite into ammonia and sulphide, respectively.

#### ABC transporters and carbohydrate-active enzymes

The genomes of cave Ktedonobacteria possess genes encoding ABC transporters belonging to several classes involved in the uptake. These include transporters for i) mineral ions, such as tungstate/molybdate (WptABC), nickel/cobalt (CbiNMQO), iron in the form of heme (CcmABCD), and phosphate (PstABCS); ii) branched-chain amino acids (LivKMHGF) and peptides (OppABCDF); iii) mono- and oligosaccharides, like alpha-glucosides (AlgEFGK), trehalose/maltose (ThuEGFK), rhamnose (RhaSPQT) and ribose/D-xylose (RbsBCAD). With respect to the uptake of mono- and oligosaccharides, cave Ktedonobacteria possess a wide array of genes encoding carbohydrate-active enzymes (CAZy) which catalyze the hydrolysis of di- and oligosaccharides deriving from complex polysaccharides such as chitin, cellulose, and hemicellulose (Table S3). However, CAZy with a signal peptide were only detected in a few MAGs from Warren cave and Imawarì Yeutà cave, suggesting that most polysaccharide-degrading enzymes are likely intracellular rather than secreted into the extracellular space. This pattern suggests that cave Ktedonobacteria are not primary degraders of complex polysaccharides but instead scavenge soluble oligosaccharides released by other polysaccharide-degrading microorganisms.

### Phylogenomics and comparative genomic analysis of Ktedonobacteria

#### Ktedonobacteria are distributed across five main ecological niches

The dataset of the Ktedonobacteria from oligotrophic caves was integrated with all the available Ktedonobacteria genomes (103 in total) retrieved from the NCBI GenBank database (Accessed in February 2023; Table S1). These genomes were grouped into five categories based on their ecological niche, vegetated soil/rhizosphere (VS/R) (35 genomes), geothermal soil/volcanic area (GS/VA) (18 genomes), permafrost/Antarctic soil (P/AS) (12 genomes), acid mine drainage/polluted soil (AMD/PS) (23 genomes), and oligotrophic soil/rock (OS/R) (7 genomes). The VS/R group includes genomes from environments like meadow soil, black locust and pine trees, and fertilised topsoil. OS/R differs from VSR by its lower nutrient availability and includes MAGs reconstructed from metagenomic data of a subsurface holobiont of the plant *Barbacenia macrantha*, which colonises nutrient-poor quartzite rocky outcrops in Brazil [31]. The GS/VA group is characterized by high temperatures including genomes recovered from thermal soils located in volcanic areas and underground burning abandoned coal mines. The P/AS group comprises MAGs obtained from tundra permafrost region and polar desert biomes sampled at different depths. These regions are characterized by extreme aridity, low temperatures, and are classified as mineral soils with high contents of calcium, magnesium, potassium, and sodium [32]. The AMD/PS includes genomes retrieved from sediments and soils associated with acid mine drainages and polluted sites.

#### Cave Ktedonobacteria belong to novel genera of *Ktedonobacteraceae* family

The phylogenomic analysis (including all the 111 Ktedonobacteria genomes) showed that Ktedonobacteria is a monophyletic class, divided into three families, i.e. *Ktedonobacteraceae*, JACDGC01 and JADMIN01 (Fig. 2). In contrast with the uncharacterized families JACDGC01 and JADMIN01, which almost exclusively included MAGs from the AMD/PS and VS/R environmental groups, *Ktedonobacteraceae* genomes were found across all environment types defined in this study, including caves. Within this family, the Imawarì Yeutà MAGs clustered closely with other MAGs from caves and were assigned to yet uncharacterised genera according to the relative evolutionary divergence (RED) values computed via GTDB-tk. A fourth family could also be identified, based on its distinct branching pattern and RED value, which is represented by a single MAG recovered from VS\R.

**Figure 2.**
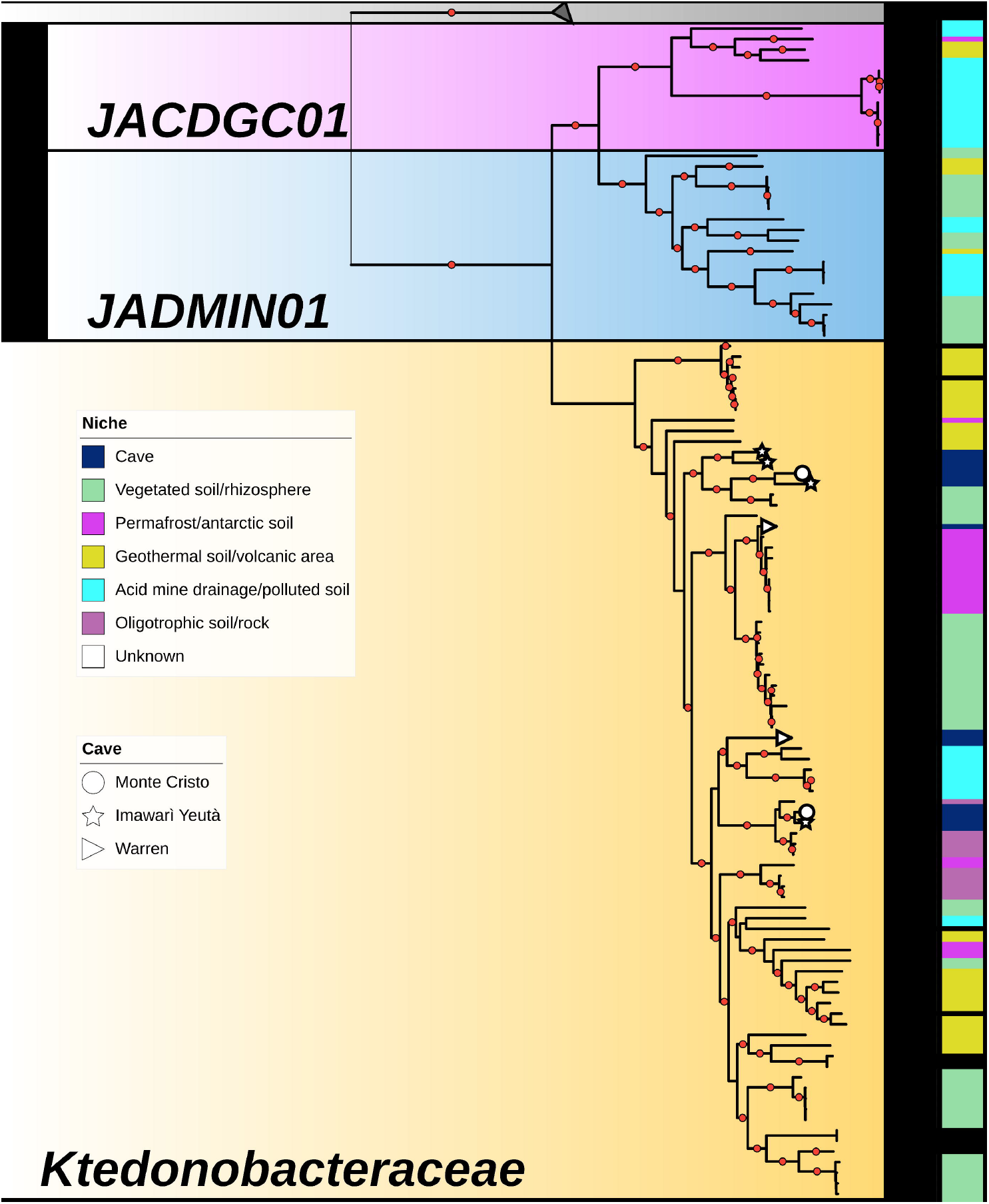
The maximum-likelihood phylogenomic tree illustrates the affiliation of the cave MAGs with the Ktedonobacteria families JACDGC01, JADMIN01, and *Ktedonobacteraceae*. Colour strips highlight the environmental niche of each genome: cave, vegetated soil/rhizosphere, permafrost/antarctic soil, geothermal soil/volcanic areas, acid mine drainage/polluted soil, oligotrophic soils/rocks, and uncharacterized habitats. Cave-specific symbols at the branch tips indicate MAGs from Monte Cristo (circle), Imawarì Yeutà (star), and Warren (triangle). Bootstrap values >80% are indicated by black circles. Collapsed clades are *Dehalococcoides mccartyi* genomes GCF_000011905.1, GCF_001889305.1 and GCF_000830885.1 that were used as outgroups (coloured in grey).

#### Taxonomy affiliation is the main driver of functional diversity in Ktedonobacteria

Comparative genomic analysis based on the functional profile of 3023 KEGG Orthologs (KO) terms revealed that both niche (R^2^ = 0.15, P = 0.001) and family (R^2^ = 0.193, P = 0.001) are significant drivers of functional variation across members of the class Ktedonobacteria (Fig. 3). However, we observed a degree of confounding between niche and family among members of JADMIN01. Indeed, most representatives of this family were retrieved from acid mine drainage environments, making it difficult to distinguish the effects of ecological niche from those of taxonomic lineage.

Of the 3023 KO terms, 919 were predicted to be family-specific (adjusted q-value < 0.001; Table S4). Among these, 67 KO terms were predominantly found in individuals of one specific family (prevalence > 0.80 in these individuals and prevalence < 0.50 in the others) (Fig. 4a). The most marked differences driven by the taxonomy lineage (different between the families) were observed in genes encoding proteins of the cell division machinery, flagellum and chemotaxis, and Type IV pili. Notably, genes of the cell division machinery were detected exclusively in *Ktedonobacteraceae*, whereas flagellar and chemotaxis genes were enriched in JACDGC01. Likewise, Type IV pili proteins were broadly present in JADMIN01 and JACDGC01 but largely absent in members of *Ktedonobacteraceae*. Other family-specific KO were genes involved in TCA cycle, sulphur metabolism, oxidative phosphorylation, energy and sulphur metabolism, carbon fixation (PPC), and atmospheric trace gas oxidation ([NiFe] hydrogenase and Cox) (Fig. 4A). Specifically, aerobic carbon monoxide dehydrogenase genes (*cox* genes) were significantly associated with the families JADMIN01 and *Ktedonobacteraceae*, while [NiFe] hydrogenase genes, which are involved in H_2_ oxidation, were almost absent in JADMIN01 and JACDGC01, while being significantly overrepresented in *Ktedonobacteraceae*.

**Figure 3.**
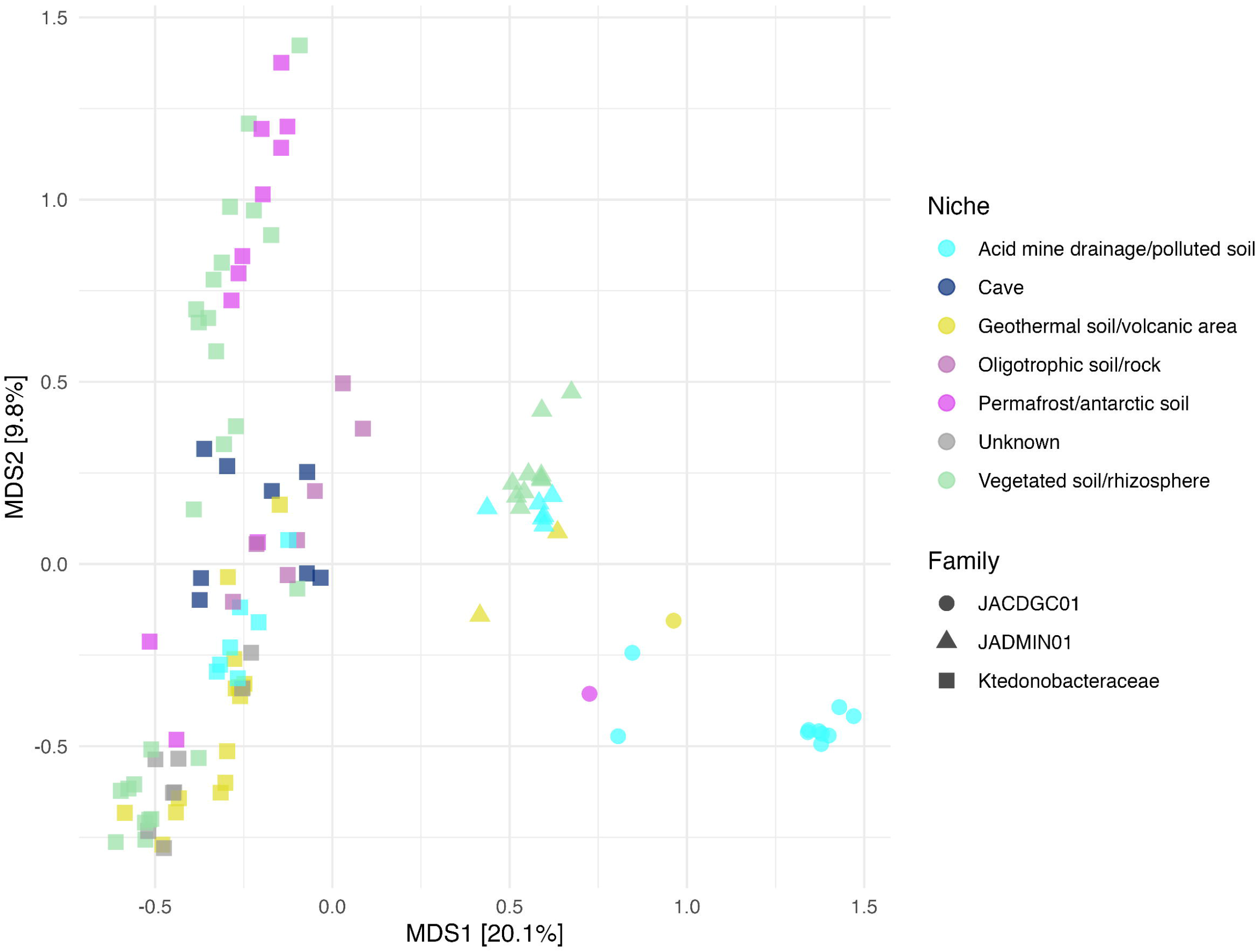
Principal Coordinates Analysis (PCoA) based on Jaccard distances computed on presence/absence profiles of KEGG orthologs across Ktedonobacteria genomes. Points are coloured according to the environmental niche of origin (acid mine drainage/polluted soil, geothermal/volcanic area, oligotrophic soil/rock, permafrost/antarctic soil, vegetated soil/rhizosphere, or unknown). Point shapes indicate taxonomic affiliation. i.e., JACDGC01 (circle), JADMIN01 (triangle), and *Ktedonobacteraceae* (square). The percentage of variation explained by each axis is shown in brackets.

**Figure 4.**
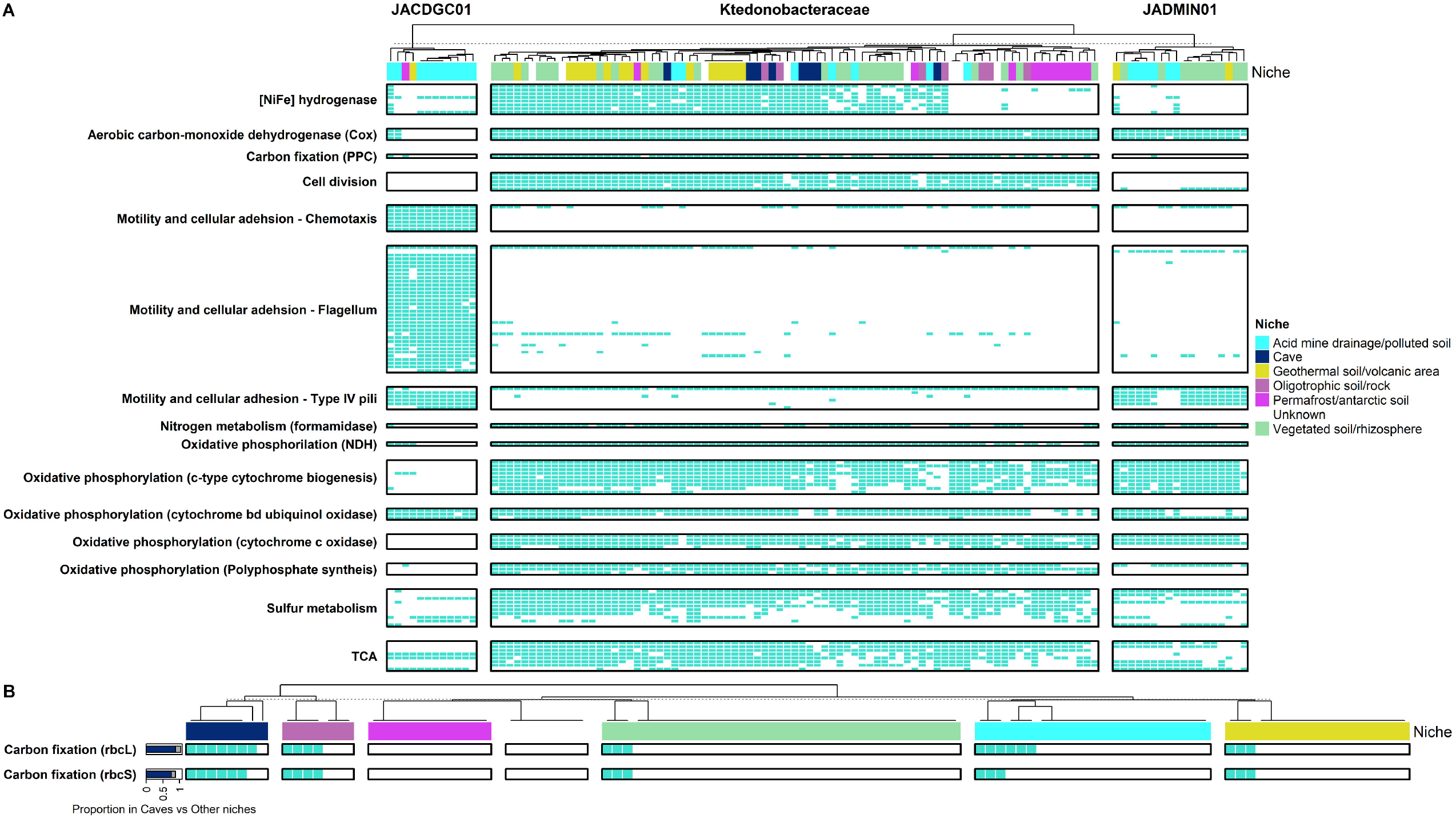
A) Heatmap showing the presence/absence of family-specific KEGG orthologs significantly associated with Ktedonobacteria families JACDGC01, Ktedonobacteraceae, and JADMIN01 (differential enrichment analysis results are available in Table S4). Columns and rows are clustered based on taxonomic groups (JACDGC01, *Ktedonobacteraceae*, and JADMIN01) and functional modules, respectively. Columns are coloured according to the environmental niche. B) Heatmap showing the presence/absence of cave-specific KEGG orthologs *rbcL* and *rbcS* genes across Ktedonobacteria genomes from different environmental niches (differential enrichment analysis results are available in Table S5). Bars show the proportion of genomes containing each gene within each environmental niche category.

#### RuBisCO genes are significantly enriched in the genomes of Ktedonobacteria inhabiting oligotrophic caves

The association between the isolation source and functional potential was also examined. Considering those functions detected in at least one MAG from each cave, a few of these were significantly enriched in Ktedonobacteria inhabiting oligotrophic cave environments (Table S5). These functions are related to toxin–antitoxin systems, lipid metabolism (e.g., soluble epoxide hydrolase), hydrolysis of carbon–sulphur compounds (L-cysteate sulfo-lyases), NAD(P)H generation (proton-translocating NAD(P)+ transhydrogenase), and the oxidation of aromatic compounds (e.g., toluene and gentisate). Moreover, key genes of the Calvin–Benson– Bassham (CBB) cycle, such as *rbcL* (ribulose-1,5-bisphosphate carboxylase/oxygenase large subunit) and *rbcS* (small subunit), were also significantly enriched (Fig. 4b). Regarding the CBB cycle, *rbcLS* genes were detected in at least one or more MAGs retrieved from each oligotrophic cave, i.e. Imawarì Yeutà, Monte Cristo and Warren caves. A few other non-cave MAGs harboured *rbcLS* genes, and they derived from oligotrophic soil/rock, geothermal soil/volcanic area, and acid mine drainage/polluted soils (Fig. 4b).

### Phylogenetic analysis of RuBisCO genes and CO_2_-fixation potential in Ktedonobacteria

#### Ktedonobacteria possess a novel Form I subtype of RuBisCO

To better understand the origin of CO_2_ fixation via CBB cycle in Ktedonobacteria, a phylogenetic tree encompassing 2,170 archaeal and bacterial RbcL proteins representing different subtypes of the Forms I, II, and III, and unknown RbcL Forms (Fig. 5a). Most Chloroflexota RbcL fall into two distantly related branches, one representing the subtype Form I’/α, considered the proto-form of RuBisCO Form I, and the other comprising the uncharacterized Form I subtype, which is phylogenetically related to Form IE. The Form I’/α branch includes Chloroflexota RbcL from the class Anaerolinae, whereas all RbcL from Ktedonobacteria cluster within the second branch together with the uncharacterized Form I subtype found in thermophilic and acidophilic Bacillota (*Kyrpidia, Acidibacillus, Sulfobacillus, Sulfoacidibacillus, Hydrogenibacillus*) and Actinomycetota. Notably, Ktedonobacteria, Bacillota, and Actinomycetota that carry the same RbcL Form I BAK (from <u>B</u>acillota <u>A</u>ctinomycetota <u>K</u>tedonobacteria) often shared overlapping geographic distributions and ecological niches (Fig. 5b). For instance, the RbcL Form I BAK was harboured by Bacillota and Ktedonobacteria colonising both acid mine drainage and volcanic/geothermal soils in Pantelleria (Italy). Likewise, Solirubrobacterales and Ktedonobacteria from oligotrophic, silica-rich soils in the Espinhaço Mountains region, where the Monte Cristo cave is located, also share the same RbcL subtype. The same pattern appears in Ktedonobacteria and Actinomycetota inhabiting MacKay Glacier and Mt. Erebus (Warren cave), both located in Victoria Land. This evidence indicates that the Form I BAK Rubisco is shared by bacteria of different lineages thriving in extreme conditions potentially conferring an advantage in nutrient-poor settings like caves and suggesting a role of horizontal gene transfer in its distribution.

**Figure 5.**
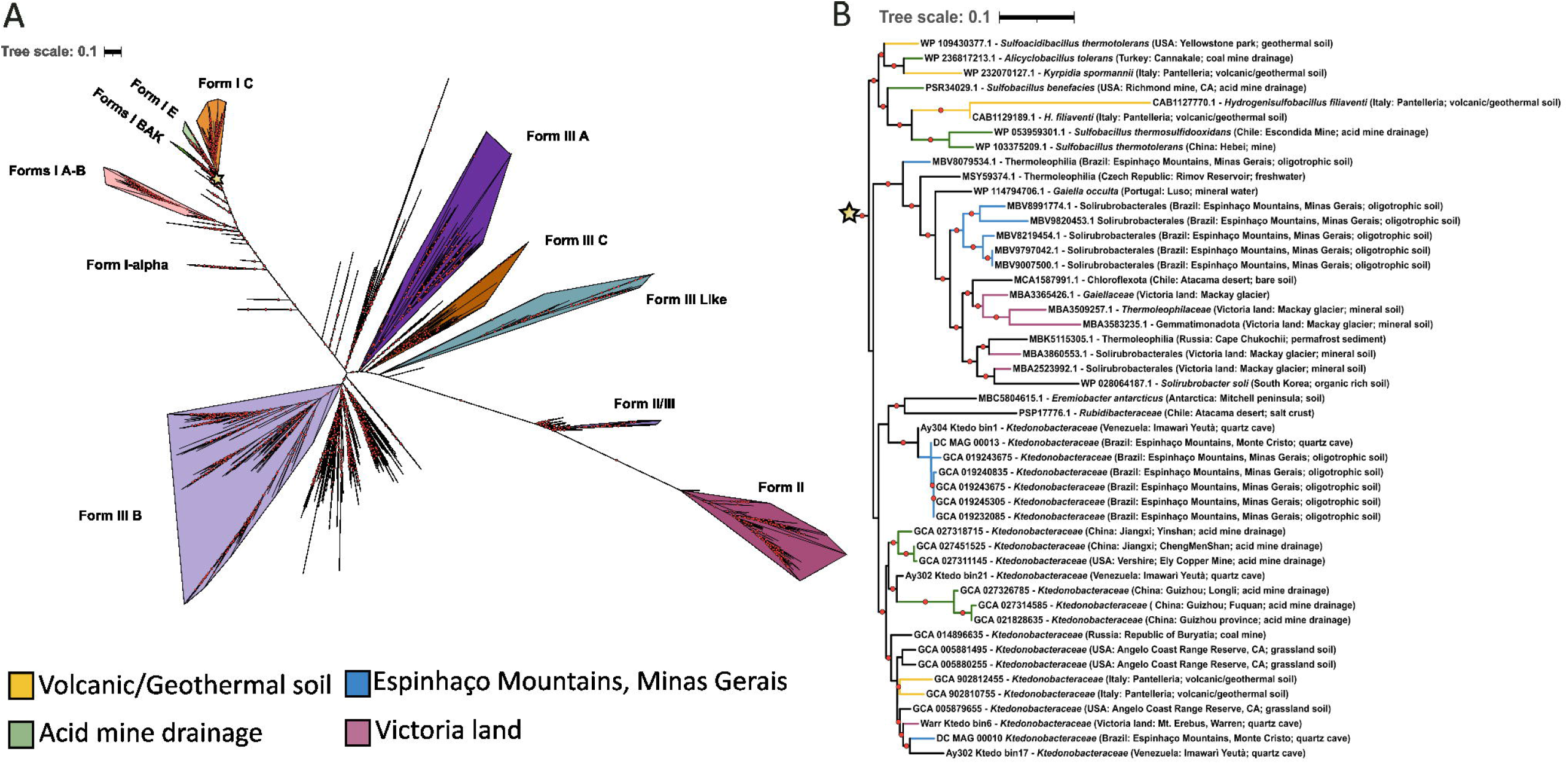
Phylogenetic analysis and environmental distribution of RuBisCO large subunit genes (*rbcL*) from Ktedonobacteria. A) Unrooted maximum-likelihood phylogenetic tree of *rbcL* representing all prokaryotic characterized and uncharacterized Forms. Clades corresponding to the different RubisCO Forms are highlighted with different colours according to Prywes et al. [24]. The interactive version of the tree shown in panel A is available at: https://itol.embl.de/shared/m11l1Qm6EqfZ: B) Expanded phylogeny showing the placement of Ktedonobacteria *rbcL* within Form I BAK (Bacillota, Actinomycetota and Ktedonobacteria). Branch colours indicate the shared environmental origin of each sequence: volcanic/geothermal soil, acid mine drainage, Espinhaço Mountains (oligotrophic soils), and Victoria Land (Antarctica). Bootstrap values >80% are indicated by black circles.

#### Ktedonobacteria possess a Form I RubisCO-mediated transaldolase variant of the CBB cycle

To further investigate the metabolic potential associated with CO_2_ fixation, Ktedonobacteria genomes were analysed for the prsecne of gene involved in regulation and functionality of the CBB cycle. This analysis revealed that the *rbc* gene cluster was conserved to some extent across Ktedonobacteria members, with some genes more conserved than others (Fig. 6). Apart from *rbcL* (K01601) and *rbcS* (K01602), other conserved genes in the cluster encoded the LysR family transcriptional regulator (*rbcR* or *cbbR*, K21703) and the red-type RuBisCO activase (*rbcX*). The *cbbR* genes were always present in two copies and always located upstream the *rbcLS* genes. The *rbc* gene cluster also included genes coding for enzymes of the CBB cycle, such as the glyceraldehyde-3-P dehydrogenase A (*gapA*, K00134) and phosphoribulokinase (*prk*, K00855).

**Figure 6.**
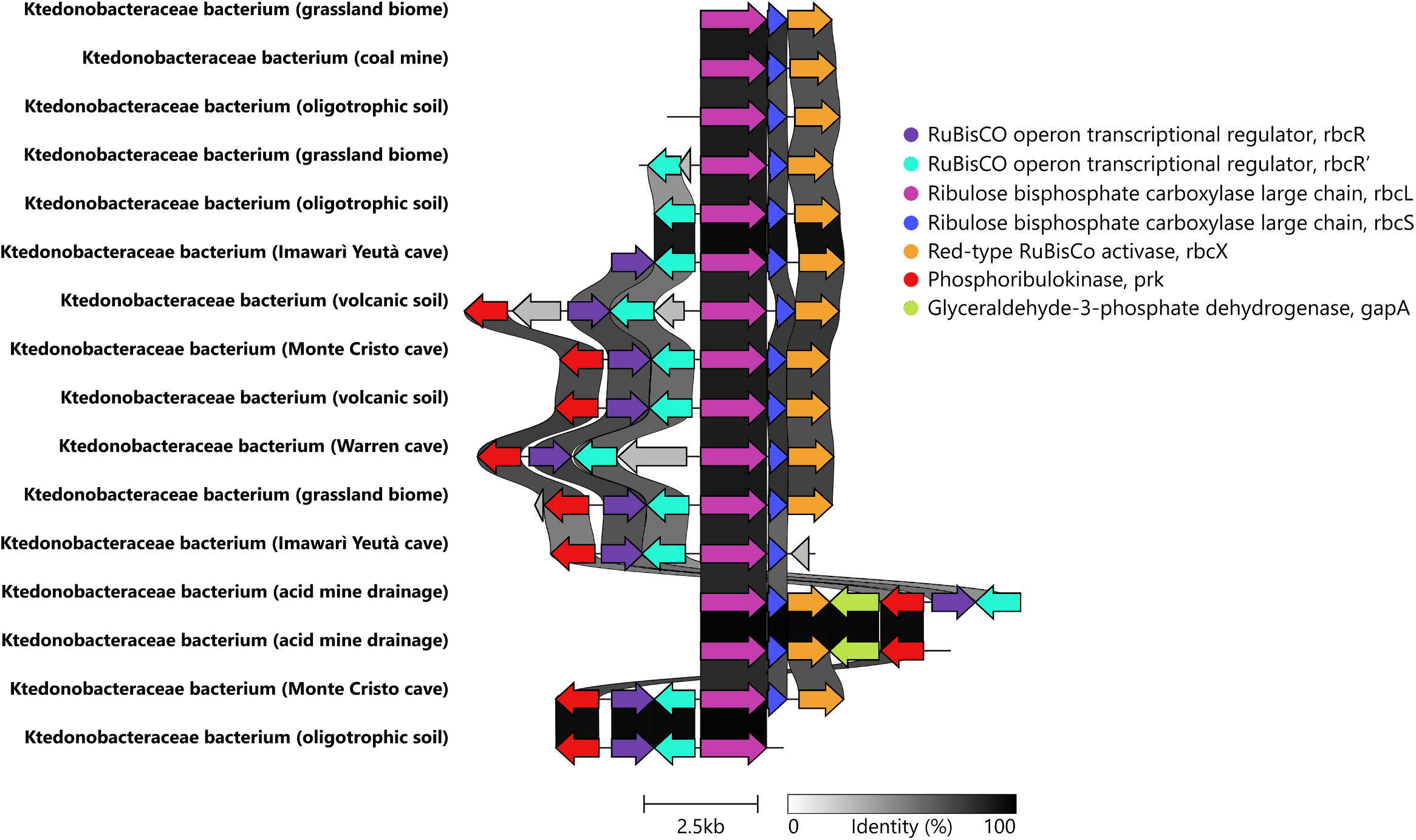
Conservation of the RuBisCO gene cluster and gene neighborhood in cave and non-cave Ktedonobacteria. Protein-coding genes are coloured according to functional annotation. Conserved syntenic blocks are highlighted by gradient bars with darker shading representing higher similarity. Genomic neighborhood comparisons are shown only for Ktedonobacteria genomes that contain a partially complete RuBiSCo cluster (*rbcR–rbcL–rbcS*).

As for the genes of CBB cycle that are not co-localized within the *rbcLS* gene cluster, we found all the genes involved in CBB cycle except for the sedoheptulose-1,7-bisphosphatase gene. Instead, we found that all the Ktedonobacteria genomes carrying at least one copy of either *rbcL* or *rbcS*, possess the genes of the transaldolase variant of the CBB cycle where the reaction of the sedoheptulose-1,7-bisphosphatase is by-passed by the enzyme transaldolase (Fig. 7). This result indicates that CO_2_ fixation in these organisms likely proceeds through an non-canonical version of the CBB cycle.

**Figure 7.**
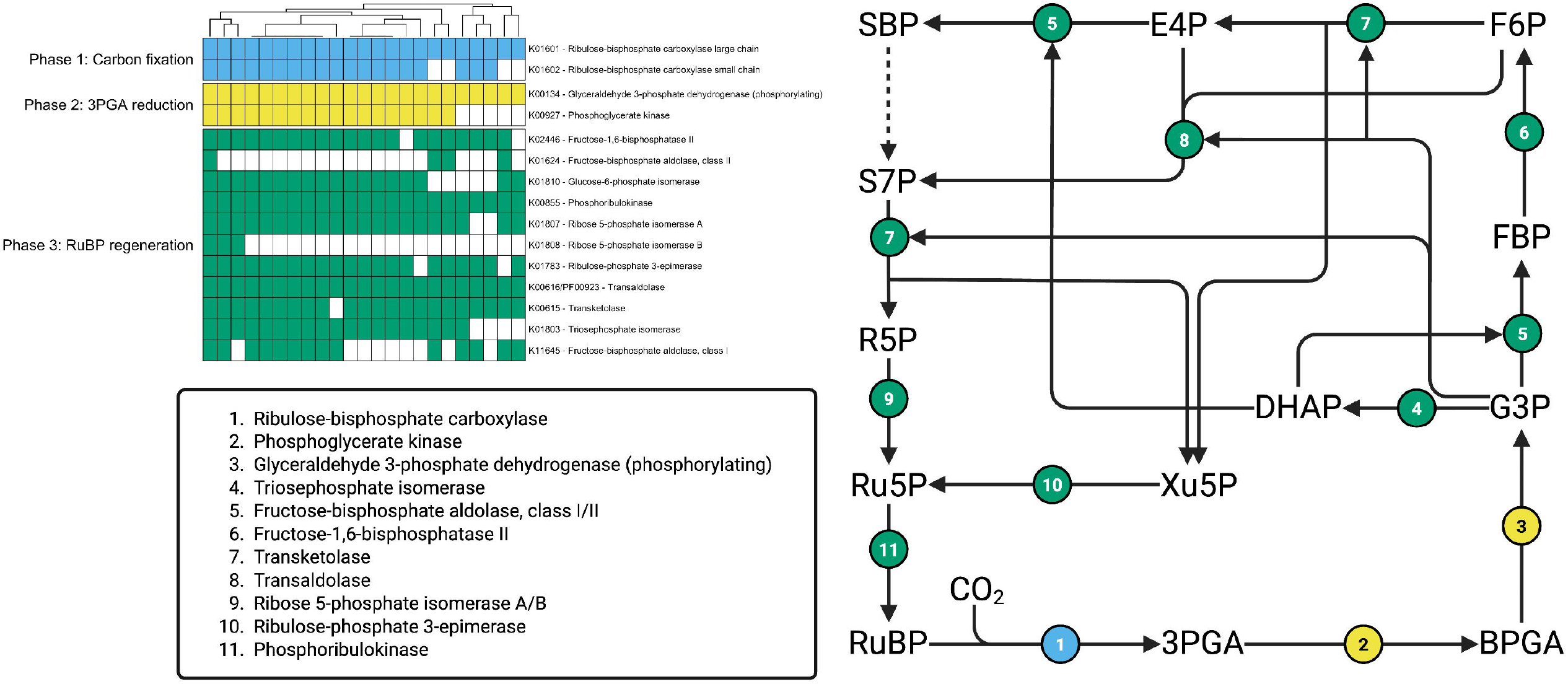
Heatmap showing the presence/absence of KEGG orthologs in Ktedonobacteria associated with the three phases of the CBB cycle, i.e., carbon fixation, 3-phosphoglycerate (3PGA) reduction, and ribulose-1,5-bisphosphate (RuBP) regeneration. Rows are clustered according to the phases of the CBB cycle (carbon fixation, 3-PGA reduction, and RuBP regeneration). In the schematic representation of the CBB cycle, coloured numbered circles map directly to the KO entries (1-11). Abbreviation: RuBP, ribulose-1,5-bisphosphate; 3PGA, 3-phosphoglycerate; BPGA, 1,3-bisphosphoglycerate; DHAP, dihydroxyacetone phosphate; G3P, glyceraldehyde-3-phosphate; FBP, fructose-1,6-bisphosphate; F6P, fructose-6-phosphate; E4P, erythrose-4-phosphate; SBP, sedoheptulose-1,7-bisphosphate; S7P, sedoheptulose-7-phosphate; R5P, ribose-5-phosphate; Ru5P, ribulose-5-phosphate.

## Discussion

In this study, we explored the metabolic potential of members of the class Ktedonobacteria, a still poorly characterised taxon within the phylum Chloroflexota, which often emerges as key players in extreme habitats characterised by nutrient limitations, extreme pH conditions and/or temperatures [33]. Among these, several studies have indicated Ktedonobacteria to be highly abundant in silica-rich oligotrophic subterranean environments such as quartzite caves in venezuelan tepuis [6, 11]. To investigate the genomic bases underlying their ability to colonize these environments, we first carried out a comprehensive analysis of the metabolic potential of Ktedonobacteria inhabiting caves, using both the MAGs reconstructed in this study and those available from a previous study. The results from phylogenomic analysis revealed that cave Ktedonobacteria belong to novel genera of *Ktedonobacteraceae* family. Furthemore, the functional analysis of the genome-based reconstructed metabolism of these Ktedonobacteria revealed the presence of functional traits associated with a mixotrophic metabolism that allows the use of both organic substrates and inorganic CO2 as carbon sources, while deriving energy from the oxidation of atmospheric trace gases or organic compounds. The ability to switch between these metabolic modes may represent a key adaptive strategy for these bacteria in oligotrophic cave environments, where the availability of organic carbon is sporadic and fluctuating [34].

The caves included in our analysis due to the occurrence of Ktedonobacteria members were Imawarì Yeutà, a quartzite cave within the Auyán Tepui in Venezuela, considered one of the oldest and most remote cave systems on Earth [9, 12, 35]; Monte Cristo Cave, developed in Proterozoic metamorphic quartzite of the San Francisco Craton in northern Brazil [29]; and Warren Cave, a fumarolic ice cave formed in volcanic rock (50–60% SiO2) [16]. Despite their distinct geological and environmental settings, these caves characterized by a high abundance of Ktedonobacteria share silica-rich substrates (>50% SiO2) in addition to strong nutrient limitation and elevated CO2 concentrations (ranging from ∼0.5% to 2% in Warren cave [36]. Conversely, Ktedonobacteria MAGs could not be detected from metagenomic data retrieved from caves opening in carbonatic or basaltic lithologies (i.e., carbonatic and sulfuric acid caves and lava tubes). This pattern may be related to the lower buffering capacity of quartz compared to carbonate substrates and to the more acidic nature of silica-based environments. This is in line with previous studies that reported Ktedonobacteria to predominate in acidic environments [33], and to decrease in their abundance in Antarctic soils as the pH increases from 5 to 7 [37]. The genomes of cave Ktedonobacteria were then incorporated into a large-scale comparative analysis including Ktedonobacteria from diverse ecological niches, in order to identify the genetic traits that distinguish cave-inhabiting lineages from those thriving in other environments. The dataset comprised all publicly available Ktedonobacteria MAGs, including genomes from geothermal soils, volcanic areas, meadows, and acid mine drainages as well as genomes from oligotrophic cave systems. The results of the differential enrichment analysis showed that the functional diversity within the class Ktedonobacteria is driven primarily by taxonomy, rather than by ecological niche. This pattern suggests that their metabolic traits are largely conserved within phylogenetic lineages, with limited influence from environmental adaptation. Similar trends have been reported for other bacterial groups with low rates of horizontal gene transfer, in which vertical inheritance of metabolic functions predominates over niche-specific gene acquisition [38]. For instance, genetic functions associated with the oxidation of atmospheric trace gases as alternative energy sources, such as genes encoding [NiFe] oxygen-insensitive hydrogenases and aerobic carbon monoxide dehydrogenases (*Cox*), were commonly found in members of the *Ktedonobacteraceae. Cox* genes were also detected in members of the JADMIN01 family, whereas hydrogenase genes were absent in both JADMIN01 and JACDGC01. The lack of these functions may account for the more limited ecological range of these two families, in contrast to the broader distribution observed for *Ktedonobacteraceae*. The ability to oxidise H_2_ enhances metabolic flexibility and confers a competitive advantage in nutrient-poor environments [39], potentially explaining the widespread occurrence of *Ktedonobacteraceae* in such habitats. Conversely, JACDGC01 is characterised by an enrichment of genes related to motility and cell adhesion, which may facilitate access to nutrient sources and promote substrate adhesion.

Although key genes involved in central carbon metabolism, nitrogen and sulphur metabolism and also atmospheric gas oxidation were correlated with taxonomy, a few functions were significantly associated with Ktedonobacteria inhabiting oligotrophic cave environments. Among these, our results highlighted the enrichment of *rbcL* and *rbcS*, which are key genes for CO_2_ fixation and primary production. In particular, our results show that *Ktedonobacteriaceae* members inhabiting subterranean environments present genetic functions involved in CO_2_ fixation through a non-canonical CO_2_ pathway known as the RuBisCO-mediated transaldolase variant. This variant of the CBB cycle was previously only identified and characterized in *Thermodesulfobium acidophilus* (of Firmicutes phylum) [30]. This chemolithoautotrophic bacterium was described to perform autotrophic CO_2_ fixation via the CBB pathway despite the absence of the sedoheptulose-1,7-bisphosphatase (SBPase) enzyme, which is typically required for the regeneration of ribulose-1,5-bisphosphate (RuBP). The bypass of the SBPase reaction is achieved through the activity of a transaldolase (TAL) enzyme, which functionally replaces SBPase by regenerating RuBP through an alternative route [30]. Notably, unlike *T. acidophilus*, which possesses a Form III RuBisCO, the *Ktedonobacteraceae* included in this study carry a Form I RuBisCO. The presence of *rbcLS* genes in members of the *Ktedonobacteraceae* from oligotrophic caves indicates that the TAL variant of the CBB cycle can also occur in association with the Form I RuBisCO, expanding the current knowledge of CBB cycle variants. This is particularly important considering that other bacterial isolates of the same family, including members of the genus *Thermogemmatispora*, possess genes associated with atmospheric gases utilisation but do not have genes encoding enzymes involved in CO_2_ fixation [40]. These findings suggest that, in Ktedonobacteria, the metabolic capacity to utilise atmospheric trace gases is evolutionarily conserved among different lineages and largely independent of habitat. In contrast, the chemolithoautotrophic potential linked to the non-canonical CBB cycle may have been acquired via HGT, as suggested by the presence of the same novel Form I subtype RuBisCO (here named BAK) found in *Ktedonobacteria* also in acidophilic members of Bacillota and Actinomycetota that share the same ecological niches. While the occurrence of chemolithoautotrophic metabolisms within the Ktedonobacteria class is a novel finding, the carbon fixation supported by the oxidation of atmospheric trace gases has previously been reported in other oligotrophic environments such as desert soils from various climatic regions [41, 42]. This process, known as atmospheric chemosynthesis, has been proposed as a major mechanisms supporting chemoautotrophic primary production in desert ecosystems where photosynthetic input is low and organic compounds are scarce [43]. However, unlike desert soils, where both atmospheric chemosynthesis and, to a lesser extent, photosynthesis contribute to primary production, deep caves lack photosynthetic activity [42]. In these ecosystems, light as energy source can be replaced by the oxidation of atmospheric trace gases and by sporadic organic carbon inputs. Consistent with this, previous studies reported a dominance of *Ktedonobacteria* during the early stages of microbial colonization in quartzite caves, not only in terms of relative abundance but also in terms of the diversity of genes associated with the oxidation of atmospheric CO and H_2_ [6]. Conversely, subsequent stages of quartzite rock microbial modification are characterized by more diverse and complex communities [6, 9, 12, 35]. Altogether, our findings provide the genomic bases outlining how Ktedonobacteria could play a key role in sustaining primary production through atmospheric chemosynthesis, initiating pioneering colonization processes that pave the way for the development of diverse microbial communities characterizing silica-rich, oligotrophic caves.

## Acknowledgements

The research on the Imawarì Yeutà cave microbiome was carried out under the speleological research permit issued by the Instituto Nacional de Parques (INPARQUES, Venezuela) and with the patronage of the Government of the Estado Bolívar, the Embassy of the Bolivarian Republic of Venezuela in Italy, and the Italian Speleological Society. We gratefully acknowledge the financial support provided by numerous private sponsors, including the Rolex Award for Enterprise, Raúl Arias with Raúl Helicópteros, Geotec S.p.A., Dolomite, Intermatica, Ferrino, Napapijri, De Walt, Scurion, Miles Beyond, and Allemano Metrology. Our sincere thanks go to the speleologists of the Theraphosa and La Venta exploring teams for their essential support during the expeditions. The authors also warmly thank Prof. Mirko Zaffagnini at UNIBO for reviewing the CBB-cycle enzymes shown in Figure 7.

## Author contributions

A.F. and M.C. designed the study, the data analysis strategy and wrote the manuscript. A.F. performed the metagenomic analyses, including MAG reconstruction, functional and statistical analyses, and prepared all figures with the support from G.B and G.R.. G.B. contributed to RuBisCO gene analyses and phylogeny. G.R. produced the initial metabolic reconstructions of cave Ktedonobacteria. D.G. extracted the DNA from the cave samples and supported data analyses. F.S. contributed to geological interpretation, field sampling, and geochemical analyses. M.C. coordinated and supervised all the research activities. All authors approved the final manuscript.

## Code and data availability

Illumina and Oxford Nanopore Technologies (ONT) sequencing reads from samples Ay302 and Ay304 are available in NCBI under BioProject accession PRJNA610757. Additional datasets deposited in Figshare (DOI: 10.6084/m9.figshare.30657227) are (i) genomes and metagenome-assembled genomes (MAGs) included in the pangenome analysis; (ii) raw data and R scripts used to reproduce Figures 3, 4, and 7 as well as the PERMANOVA analyses; (iii) Anvi’o genome and pangenome databases, including the pangenome summary file; and (iv) IQTREE input and output files used for the *rbcL* phylogenetic reconstruction (in Figure 5).

