## Supplementary Tables for "Genomic evidence for atmospheric chemosynthesis as a defining trait of Ktedonobacteria inhabiting subterranean environments": Table S2.docx

**Table S2**. Description of and relative abundance of Ktedonobacteria in the oligotrophic cave samples included in this study

| **Cave** | **Sample name** | **Description** | **Mineralogy** | **SiO_2_ (%)** | **pH** | **Temperature**  **(° C)** | **Abundance of Ktedonobacteria**  **(%)** | **Reference(s)** |
| --- | --- | --- | --- | --- | --- | --- | --- | --- |
| Imawarì Yeutà | Ay304 | White dendritic biofilm on a quartzite rock on the cave floor | Quartz (SiO_2_) | >98 | 5 | 14.9 | 67 (Illumina MiSeq ^c^) | [6, 12] |
| Imawarì Yeutà | Ay317 | Pristine quartzite cave wall | Quartz (SiO_2_) | >98 | 4 | 14.9 | ~5 (Illumina MiSeq) | [9, 12] |
| Imawarì Yeutà | Ay302 | White part of the amorphous silica speleothem on a quartzite rock wall | Quartz, Opal-G (SiO_2_) | >98 | 5 | 14.9 | ~5 (Illumina MiSeq) | [9, 12] |
| Warren | - | Sandy sediments | Anorthoclase feldspar [(Na,K)AlSi_3_O_8_)] | 56 | 5.2 | 14.6 | 63 (clone library ^d^) | [16] |
| Monte Cristo | P7 | Wet brownish saprolite (weathered rock) | Quartz (SiO_2_), muscovite  [KAl_2_(Si_3_Al)O_10_(OH, F)_2_],  kaolinite [Al_2_Si_2_O_5_(OH)_4_],  rutile (TiO_2_) | >90^a^ | 5.5-7.1 | ~20.0^b^ | 10^e^ | [29] |

^a^ Typical % of SiO_2_ in a quartzite cave

^b^ Mean annual temperature of Diamantina, Minas Gerais (Brazil)

^c^ Relative abundance based on Illumina MiSeq 16S rRNA gene sequencing data

^d^ Relative abundance based on 16S rRNA gene clone library analysis

^e^ % of MAGs attributed to Ktedonobacteria (2 MAGs over 20 total numebr of MAGs)
